# Differential modulation of mouse heart gene expression by infection with two *Trypanosoma cruzi* strains: a transcriptome analysis

**DOI:** 10.1101/574343

**Authors:** TBR Castro, MCC Canesso, M Boroni, DF Chame, D de Laet Souza, NE Toledo, EB Tahara, SD Pena, CR Machado, E Chiari, AM Macedo, GR Franco

## Abstract

The protozoan *Trypanosoma cruzi* (*T. cruzi*) is a well-adapted parasite to mammalian hosts and the pathogen of Chagas disease in humans. As both host and *T. cruzi* are highly genetically diverse, many variables come into play during infection, making disease outcomes difficult to predict. One important challenge in the field of Chagas disease research is determining the main factors leading to parasite establishment in the chronic stage in some organs, mainly the heart and/or digestive system. Our group previously showed that distinct strains of *T. cruzi* (JG and Col1.7G2) acquired differential tissue distribution in the chronic stage in dually-infected BALB/c mice. To investigate changes in the host triggered by the two distinct *T. cruzi* strains, we assessed the gene expression profile of BALB/c mouse hearts infected with either JG, Col1.7G2 or an equivalent mixture of both parasites during the initial phase of infection. This study demonstrates a clear distinction in host gene expression modulation by both parasites. Col1.7G2 strongly activated Th1-polarized immune signature genes, whereas JG showed only minor activation of the host immune response. Moreover, JG strongly reduced the expression of genes for ribosomal proteins and mitochondrial proteins related to the electron transport chain. Interestingly, evaluation of gene expression in mice inoculated with the mixture of parasites showed expression profiles for both up- and down-regulated genes, indicating the coexistence of both parasite strains in the heart during the acute phase. This study suggests that different strains of *T. cruzi* may be distinguished by their efficiency in activating the immune system, modulating host energy and reactive oxygen species production and decreasing protein synthesis during early infection, which may be crucial in defining parasite persistence in specific organs.

**Author Summary:** The causative agent of Chagas disease, *Trypanosoma cruzi*, retains high genetic diversity, and its populations vary greatly across geographic locations. The *T. cruzi* mammalian hosts, including humans, also have high genetic variation, making it difficult to predict the disease outcome. Accordingly, this variability must be taken into account in several studies aiming to interrogate the effect of polyparasitism in drug trials, vaccines, diagnosis or basic research. Therefore, there is a growing need to consider the interaction between the pathogen and the host immune system in mixed infections. In the present work, we present an in-depth analysis of the gene expression of hearts from BALB/c mice infected with Col1.7G2 and JG alone or a mixture of both strains. Col1.7G2 induced a higher Th1 inflammatory response, while JG exhibited a weaker activation of immune response genes. Furthermore, JG-infected mice showed a notable reduction in the expression of genes responsible for mitochondrial oxidative phosphorylation and protein synthesis. Interestingly, the mixture-infected group displayed changes in gene expression as caused by both strains. Overall, we provided new insights into the host-pathogen interaction in the context of single and dual infection, showing remarkable differences in host gene expression modulation by two *T. cruzi* strains.

## Introduction

Chagas disease (CD) is a parasitic illness caused by the kinetoplastid protozoan *Trypanosoma cruzi*. Six to seven million people are estimated to be compromised by this disease, which affects mostly poor communities in rural areas of Latin America. [1]. Despite being the prototype of a neglected tropical disease, CD has recently gained attention in nonendemic areas due to increasing emigration of affected people from endemic to nonendemic countries, with new cases occurring mainly by infected blood transfusion, organ transplantation and congenital transmission [2–4]. Currently, there are six discrete typing units (DTUs I-VI) described for *T. cruzi*, according to a series of genetic markers, such as rDNA 24Sα, miniexon and mitochondrial polymorphisms [5–8], and a seventh has been postulated (Tcbat) [9]. This broad genetic diversity makes *T. cruzi* a highly complex organism and also plays an essential role in the differential tropism in host tissues, which culminates in diverse clinical manifestations observed in chronic patients and experimental models of CD [10–13]. Notably, the term tropism has been used with different meanings by many authors in the scientific literature [14–16], and here, we refer to tissue tropism as the ability of a particular pathogen to infect and persist within an organ or set of organs [17].

Elucidating the molecular mechanisms dictating the interaction between the pathogen and its host is crucial for understanding the disease progression and the development of new treatments. Previously, Andrade *et al.* showed that after inoculating BALB/c mice with a mixture of different strains of *T. cruzi*, namely, JG (*T. cruzi* I) and Col1.7G2 (*T. cruzi* II), the two parasite strains did not evenly distribute among different tissues in the chronic phase of the disease. JG was primarily found in the heart, while Col1.7G2 was encountered in the rectum of the animals [11]. Curiously, mice infected with only one strain did not exhibit this pronounced tissue tropism. In addition, different mouse lineages such as C57BL/6J and SWISS were unable to reproduce the aforementioned tissue tropism, indicating the role of the host in the different behaviors of *T. cruzi* strains [12]. In parallel, such tropism was also detected in human patients during the chronic phase of CD, as different organs were the subject of distinct *T. cruzi* DTUs establishment, leading to the proposition of the ‘clonal-histotropic model’ hypothesis [10, 11, 18].

Recently, extensive research has helped to better understand the immunological and molecular interaction between *T. cruzi* and its mammalian host. An efficient and non-exaggerated immune response is crucial to pathogen clearance, without much damage to the host tissue [19]. The innate immune system provides the first line of defense to initiate an effective response against a parasite via pattern recognition receptors (PRRs), of which the Toll-like receptors (TLRs) are the best known [20]. Among the most well-studied TLRs in the context of CD are *Tlr-2* and *Tlr-9* [21]. The Glycosylphosphatidylinositol (GPI)-anchored mucin-like glycoproteins (tGPI-mucin), widely present on the parasite surface, and unmethylated CpG DNA sequences are the primary immunostimulatory ligands of these TLRs, respectively. They have been recognized to trigger the release of *Il-12* and *TNF* by dendritic cells and macrophages, which are pivotal for host resistance at the beginning of the acute phase of CD [22, 23]. Moreover, the establishment of a Th1 response, which is dependent on the release of *Ifn-γ*, has been extensively studied and is essential for parasite control and host survival [21, 24]. However, there is a lack of comparative studies showing the peculiarities of the host response against distinct strains of *T. cruzi* and their role in differential parasite tissue preferences. Experiments in mice and rats comparing JG (*T. cruzi* II) and CL-Brenner (*T. cruzi* VI) have shown remarkable differences in the systemic production of inflammatory cytokines and inflammatory cells [25, 26]. Even though much has been achieved to understand the complexity of tissue preferences in the parasite, real changes in host gene expression during infection with distinct *T. cruzi* strains and mixtures have not yet been investigated.

In the present work, we report a thorough gene expression analysis of BALB/c hearts during the acute stage of infection by different strains of *T. cruzi* and their mixture. Our data reveals that JG-infected mice display less pronounced induction of both innate and adaptive immune response genes in contrast to Col1.7G2-infected mice. Moreover, biological processes such as translation and mitochondrial oxidative phosphorylation are intensely downregulated in JG, indicating a cellular metabolism modulation that might benefit JG parasites. Remarkably, the mixture-infected animals showed both profiles simultaneously. Our data demonstrate the complexity of host-pathogen interactions in the context of experimental CD and how distinct *T. cruzi* strains differently affect gene expression in mouse hearts.

## Results and Discussion

### Differential performance of infection by JG, Col1.7G2 and their mixture in BALB/c mice

One critical aspect of distinct *T. cruzi* strains is their high genetic variability and differential parasitism when infecting a mammal host. Our research group has intensively studied the Col1.7G2 (*T. cruzi I*) and JG (*T. cruzi II*) strains regarding characteristics of their infection in different mouse lineages and tissue tropism [12, 13, 27]. For example, the JG strain rarely causes death in BALB/c mice, while Col1.7G2 presents high virulence, killing these animals during early moments of infection (Fig 1A). On the other hand, this virulence does not appear to be caused by parasite burden, since JG display higher parasitemia than Col1.7G2 in the acute phase of our experiments (Fig 1B). Interestingly, animals infected by an equivalent mixture of both strains present high parasitemia, and a high mortality rate (Fig 1A-B). Such phenomenon is also noted by Campos *et al.*, in which BALB/c mice infected by two *T. cruzi* I strains (AQ1-7 and MUTUM) present undetectable parasitemia, but when coinfected with JG, a synergic effect in parasitemia is observed [28]. In general, highly multiplicative intracellular pathogens exacerbate the immune response and promote early activation of CD8+ T cells [29]. This effect does not appear to be the case of JG, since experiments confirmed the presence of lower levels of inflammatory cytokines in infected mice and rats during the acute phase in comparative studies [25, 26]. A possible explanation for this diverse mortality and parasitemia levels may be the exacerbation of the inflammatory response by virulent strains, and the remarkable ability to evade the innate mechanism presented by nonvirulent strains.

**Figure 1.**
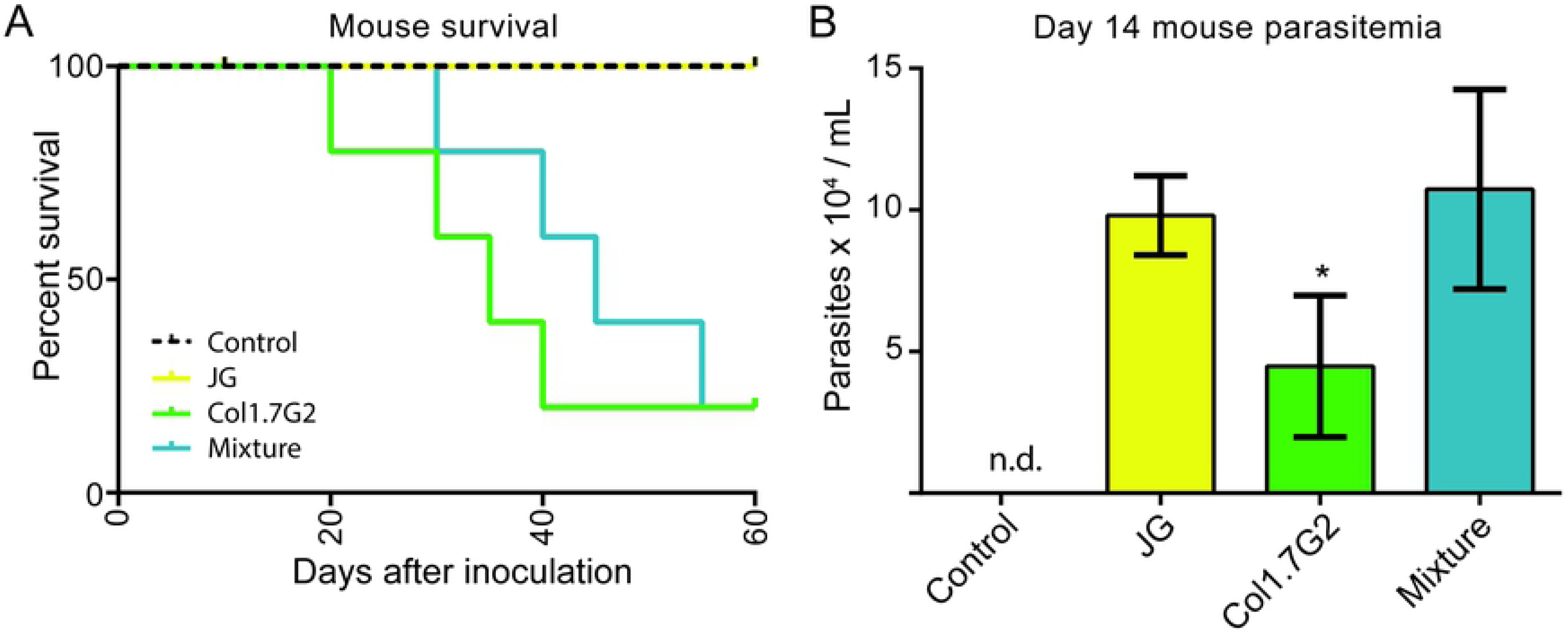
Differences between two *T. cruzi* strains. (A) Survival curve of mice with *T. cruzi* infection for 60 days. (B) Mouse parasitemia 14 days after infection. Data represents mean ± SEM, n=5 mice/group * p<0.05 comparing Col1.7G2 group to JG and mixture.

Currently, the reason for strain preferences for particular tissues is not completely understood. In humans, it has been suggested that distinct *T. cruzi* strains can prevail in different organs and cause variable damage to its hosts, culminating in distinct CD outcomes in the chronic phase [10, 11]. Thus, different forms of CD would be dependent on the parasite strain and its tropism for specific niches and on the host genetic variability, a concept from which the ‘clonal-histotropic model’ for the pathogenesis of the disease was constructed [30]. Severe chronic manifestations of CD result from persistence of the intracellular stage of the parasite (amastigotes) in heart and muscle cells, enteric ganglia, and adipose tissue [31, 32]. It is still a matter of controversy whether *T. cruzi* parasites exhibit true tissue tropism for heart cells due to tissue-specific protein-protein interactions in the cell surface, or if differential proliferation and selection by immune system/drug resistance affect survival and persistence, leading to the outcome observed in chronic stages [33]. At the acute stage of infection, *T. cruzi* parasites can infect virtually any nuclear cell or organ, bringing the real meaning of the tissue tropism concept to question. We cannot discard the hypothesis that the differential strain ‘preference’ may represent the consequence of competition between two or more distinct *T. cruzi* strains for a specific niche, as determined by strain resistance and adaptation to the immune system. Experiments with Holtzman rats that attested to the efficiency of the JG strain to persist in the heart, compared to the CL-Brener strain (*T. cruzi* VI), at the end of the acute phase as well as the chronic phase [26]. It has been suggested that *T. cruzi* persists preferentially in cells with high rates of fatty acid metabolism, and therefore, those cells can provide adequate nutrients for the parasites [34]. However, it is interesting to note that cells where *T. cruzi* parasites are predominant in the chronic phase, such as adipocytes, cardiomyocytes, skeletal fibers, and neurons, are the cells with minor turnover rates in mammals [35] and may represent tissues that can harbor latent parasites for long periods.

### Heart transcriptome acquisition, quality assessment, and gene clustering

To help us to understand the modulation of heart gene expression by parasite infection, we infected BALB/c mice with the JG, Col1.7G2 and an equal mixture of both strains (Fig 2A). mRNA from independent biological replicates of each group was sequenced to generate up to 120 million paired-end reads in total (S1 Table), followed by processing through our RNA-Seq analysis pipeline (S1 Fig). The tendency within our data was evaluated by executing principal component analysis (PCA) using the regularized-log (log2) transformed data matrix from normalized read counts. The dimensionality reduction through PCA evidenced the separation between control and experimental groups by the first principal component. The second principal component separated Col1.7G2 and JG-infected groups, while the mixed infection were distributed between both profiles (Fig 2B). The relationship among samples was revealed using a heatmap of the Euclidean distances between samples. As seen before, experimental and control groups clustered separately, and Col1.7G2 and JG groups are shown to be distinctive. Interestingly, the mixture group shows moderate similarity among samples from each infected mouse, but the control group does not (Fig 2C).

**Figure 2.**
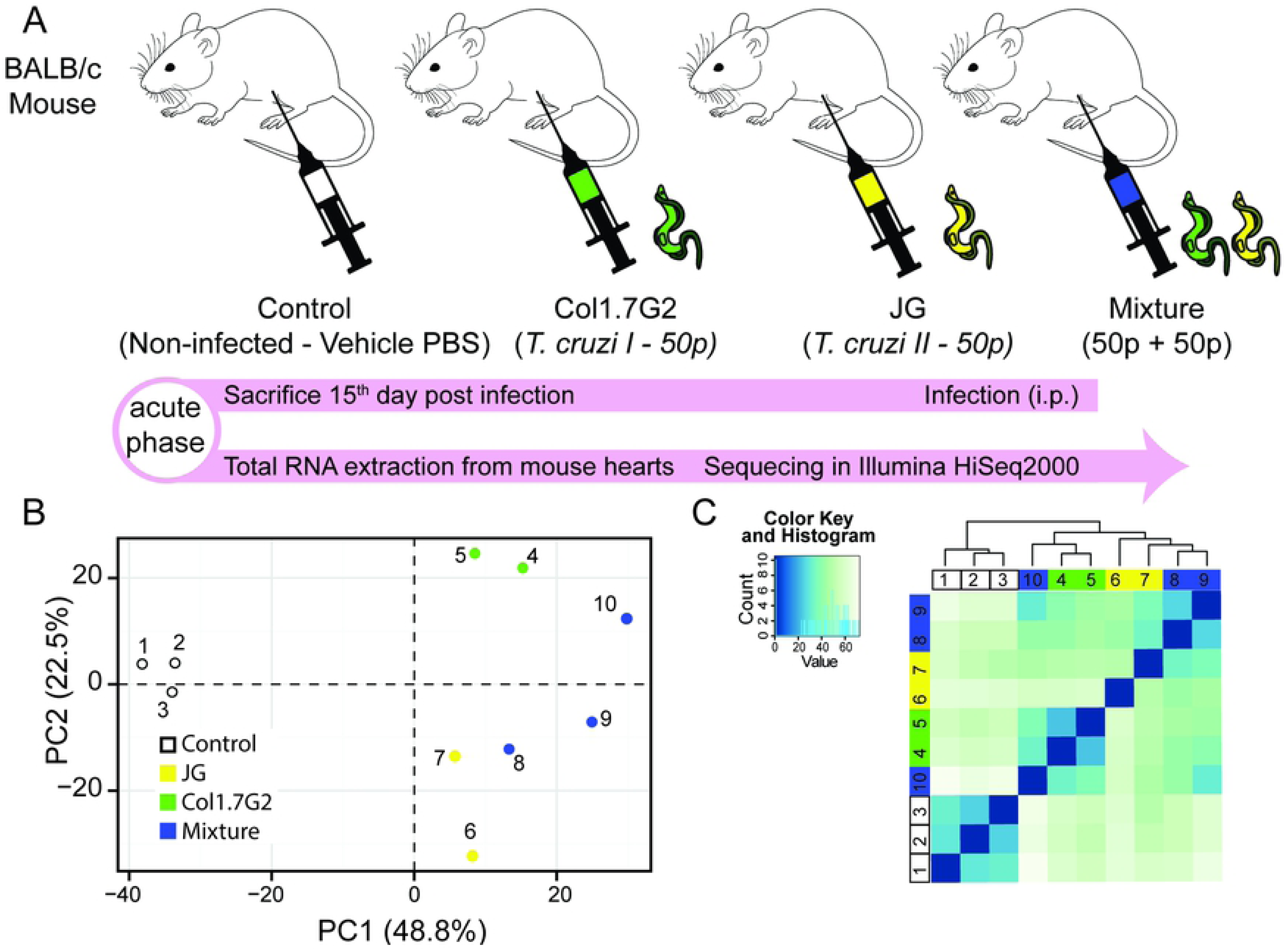
Experimental Design. (A) BALB/c mice were divided into one control and three experimental groups, infected with Col1.7G2, JG, or mixed strains of equivalent quantities of both strains. After 15 days, during the acute phase, mice were euthanized, and hearts were removed for total RNA extraction and sequencing with Illumina HiSeq2000. (B) To capture overall expression profile of each infected mouse sample, controls (blank dots), Col1.7G2-infected (green dots), JG-infected (yellow dots) and Mixture-infected (blue dots), were compared using principal component analysis (PCA). The first two principal components (PC1 and PC2) are plotted according to rlog-transformed data of read counts for each gene. (C) Hierarchical clustering of rlog-transformed data matrix exhibited as heatmap of Euclidian distances between samples.

### Revealing the Differentially Expressed Genes (DEGs)

We found approximately 16,400 genes expressed in mouse hearts in all experimental groups (S2 Table). 2,583 (S3 Table) genes presented a minimum 2-fold change (1 log2 Fold Change) with a 1% false discovery rate (FDR) when comparing infected groups to the control non-infected groups and were considered differentially expressed genes DEGs. From these, 2,396 genes are protein coding, 47 are long intervening noncoding RNA (lincRNA), and 140 belong to other biotypes (S4 Table). The distribution of DEGs in each group is represented by volcano plots (Fig 3A-C). It is worth noting that there are remarkable differences in fold changes of DEGs among groups. Col1.7G2-infected animals exhibit more upregulated (999 in Col1.7G2 and 729 in JG) and less downregulated DEGs (145 in Col1.7G2 and 673 in JG) relative to the control group in comparison to JG infected-animals. Notably, mouse hearts from the mixed infection presented a higher number of downregulated DEGs (775) and a higher number of upregulated DEGs (1278) relative to the control group, compared to both single-infected groups.

**Figure 3.**
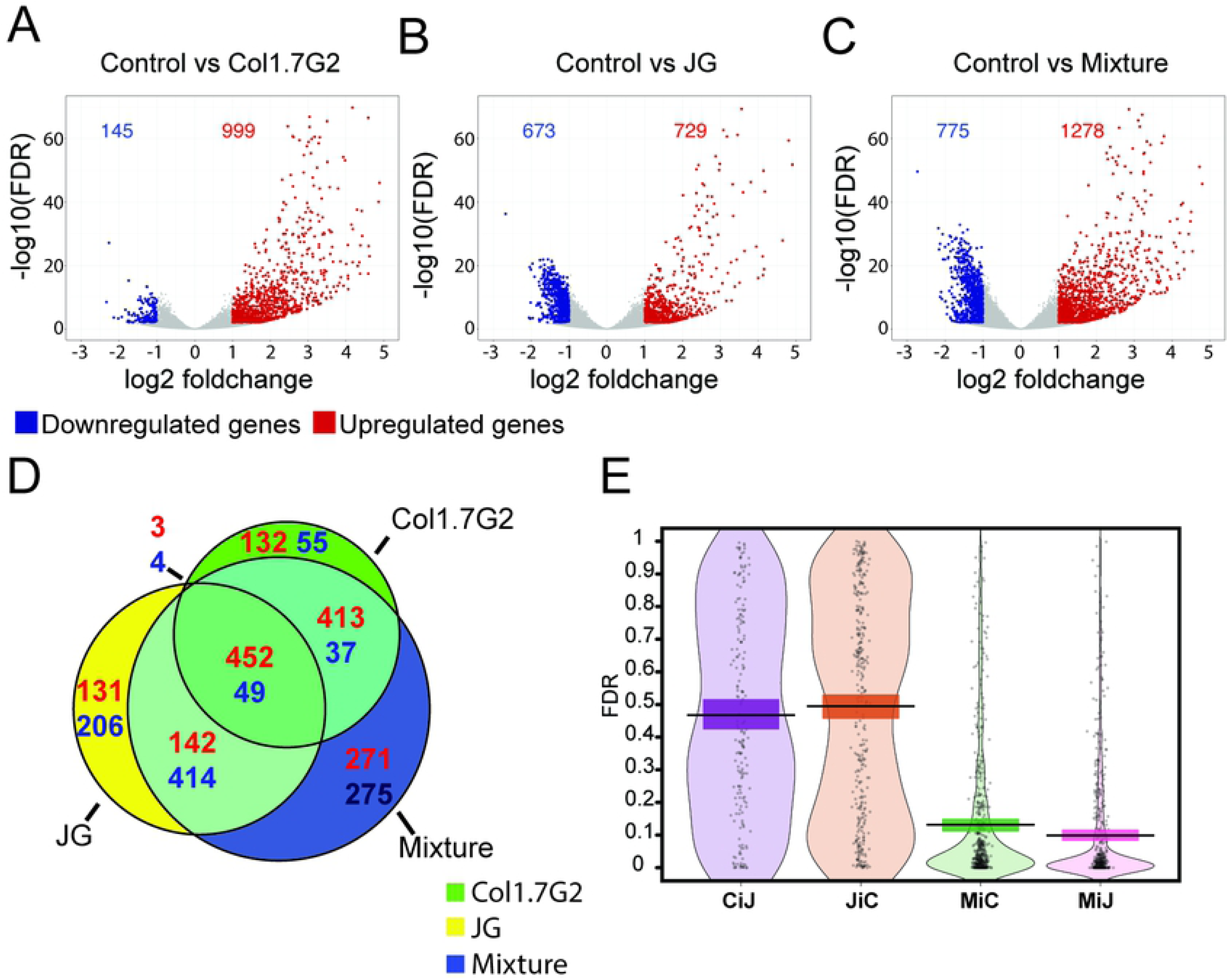
Comparative analysis of mouse gene expression after exposure to single and mixed *T. cruzi* infections. Volcano plot of (A) Col1.7G2-, (B) JG- and (C) mixture-infected groups showing log2 of fold changes of each detected gene on the x-axis, along with -log10 of adjusted p-values (FDR) after comparison with noninfected controls. Differentially expressed gene are in blue (log2fold-change < -1) for downregulated genes and red (log2fold-change > +1) for upregulated genes. Grey dots represent genes that were not considered differentially expressed. Genes with -log10 FDR equal or smaller than 50 are shown as diamond shapes. (D) The scaled Venn diagram shows the proportion of shared and exclusive genes in each group. Up- and downregulated genes are shown in red and blue, respectively. (E) Distribution of FDRs of exclusive genes from mixture in the Col1.7G2-infected (MiC) or JG-infected (MiJ) groups and exclusive genes from Col1.7G2 in JG-infected group (CiJ) or vice versa (JiC) dataset are shown.

At the acute stage of the mixed infection, it is improbable that many host cells are infected by both strains, since we inoculated mice with a very low load of parasites in both single and mixed infections. Therefore, based on the observation that in the acute phase of mixed infection, hearts present both JG and Col1.7G2 expression patterns, but in the chronic phase, Col1.7G2 seems to be eliminated from this organ, as seen by Andrade *et al*., 1999, we suggest that JG proliferates at a higher rate than Col1.7G2, predominating in the heart in the chronic phase. This hypothesis corroborates previous reports showing that both JG and Col1.7G2 were detected in the hearts of dually-infected BALB/c mice by low-stringency PCR in the acute phase of infection, but at the chronic stage (3 to 6 months after infection), there was a predominance of JG in the hearts of these animals [12, 13]. Additionally, recent findings by Dias *et al*. demonstrated that Col1.7G2 is less efficient than JG in proliferating in the cardiomyocytes of neonate BALB/c mice in culture, which reinforces our hypothesis [36].

We asked if the transcriptome profile of infected mice would differ between *T. cruzi* strains. Thus, we investigated whether DEGs are shared or exclusively expressed in a specific experimental group, as shown in a Venn diagram (Fig 3D). A total of 501 (25%) DEGs are shared among all groups and can represent genes intrinsically influenced by the *T. cruzi* infection (S5 Table). Interestingly, for exclusive DEGs, the mixture-infected animals show an equivalent number of up- and downregulated genes (271 up and 275 down), while JG-infected animals present more downregulated genes (132 up and 206 down), and Col1.7G2- present more upregulated (131 up and 55 down) DEGs. Most DEGs from the Mixture-infected group (1507 or 74%) correspond to DEGs also found in JG- and Col1.7G2- single infected animals. We decided to analyze the exclusive DEGs from the mixture-infected group (∼21%), since it remains an open question whether a mixed infection leads to exacerbation of the pathological effect of single strains (synergistic effect) or if it is merely the sum of the impact of both strains [25, 37, 38]. We concluded that the mixed infection is much more similar to the sum of every infection than previously understood. We examined the FDR distribution of exclusive DEGs from Col1.7G2, JG and Mixture-infected groups that were not considered as DEGs in the other groups (Fig 3E). Notably, an accumulation of mixture-exclusive DEGs close to the threshold to also be considered a DEG (FDR < 0.01) in the JG or Col1.7G2 groups, but this tendency was not observed for exclusive JG DEGs present in the Col1.7G2 group and vice-versa, demonstrating that these two groups are distinct. Thus, it is plausible that these two parasite strains induce different gene expression responses in the host, and that mixture-infected animals combine cells infected by both strains. The effect of gene expression modulation in the heart of mixture-infected animals will depend on how many cells are infected by JG or Col1.7G2, leading to profiles that are more similar to Col1.7G2 single infection or to JG single infection.

### Different *T. cruzi* strains stimulate distinct gene expression response in mouse hearts

We next asked which biological processes are mainly represented by DEGs, and whether differences could be detected in the more representative categories when comparing the different groups. To achieve this, we separately analyzed the genes upregulated and downregulated upon JG, Col1.7G2, or mixture infection versus uninfected control in mouse hearts for functionally enriched biological processes (GO - Gene Ontology analysis) (Fig. 4 and S6 Table). For upregulated genes, we found that all groups present significantly enriched biological processes related to the innate, adaptive immune, and inflammatory responses of the host against the pathogen (among the top 30 most represented GO categories in these groups – Table S6). We also found significantly enriched categories of cellular response to interferon-beta and gamma, and defense response to virus, bacteria, and protozoa. This result highlights the intersection of intracellular pathogen defense mechanisms. However, the number of DEGs was not similar in each category. Col1.7G2-infected and mixture-infected groups displayed the highest number of DEGs in all upregulated biological processes (Figs 4A and 4C, respectively) and the JG-infected group presented the lowest number of DEGs in the upregulated categories (Fig 4B). Notably, both the mixture and Col1.7G2-infected groups presented enriched biological processes with statistical significance (-log10 pvalue) that were higher than those encountered in the JG-infected group. This observation correlates with previous studies showing that JG leads to lower cytokine production and immune system activation during acute infection [25]. Every DEG encountered in each GO is available in the supplementary material (Table S7), as well as all non-DEGs from the background (Table S8).

**Figure 4.**
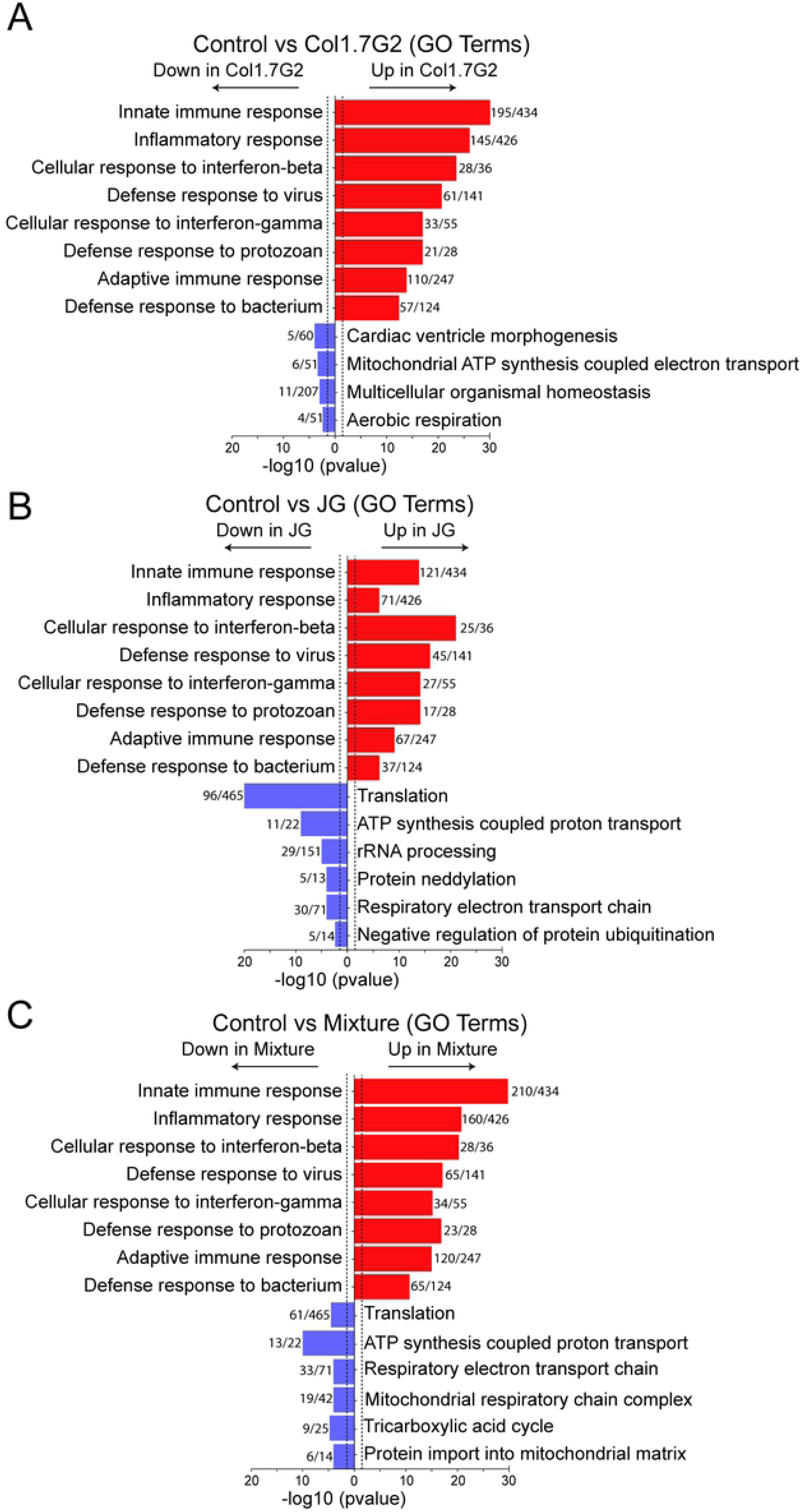
Analysis of enriched Gene Ontology categories present in differentially expressed genes. All genes from (A) Col1.7G2, (B) JG and (C) mixture groups were analyzed in accordance with their Gene Ontology terms using the topGO package for R. Encountered GO terms derived from up and downregulated genes are shown in red and blue, respectively. Bars represent -log10 of Fisher elim p-value (p < 0.01). Histogram values represent the number of differentially expressed genes by the number of expressed genes in the background.

Remarkably, the enriched biological processes observed in the mixture-infected group are consistent with those seen in the individual infected groups (Fig 4C). For instance, a high number of upregulated genes were related to immune response (Col1.7G2 profile), and a high number of downregulated genes were related to translation, ATP synthesis and respiration (JG profile). Hence, the coexistence of both profiles in the mixture is substantial evidence of the presence of both *T. cruzi* strains in the mice during the acute phase of infection. Both JG- and the mixture-infected groups displayed categories of protein translation, ATP synthesis coupled respiratory electron chain and a significantly enriched tricarboxylic acid cycle.

To visualize the interactions between the enriched pathways and the genes within our list of DEGs, we constructed a functional network of each infected group compared to the control using the Cytoscape plugin, ClueGO. Here, we can appreciate the Col1.7G2 activated infection genes (Fig 5A). Protein phosphorylation mediated by MAPKinase follows the transcription factor NfkB activation with subsequent immune-mediated pathway stimulation, such as cytokin production that leads to the establishment of a strong immune response [39]. Interestingly, JG infected samples exhibit less immune-mediated pathways when compared to Col1.7G2. However, it is evident that mitochondrial processes and electron transport chain genes are downregulated due to JG infection, particularly during the early phase (Fig 5B). It is worth noting that the Mixture-infected group displays a rich immunological network and also mitochondrial related gene downregulation, evidencing the concurrently presence of Col1.7G2 and JG-infected cells (Fig 5C).

**Figure 5.**
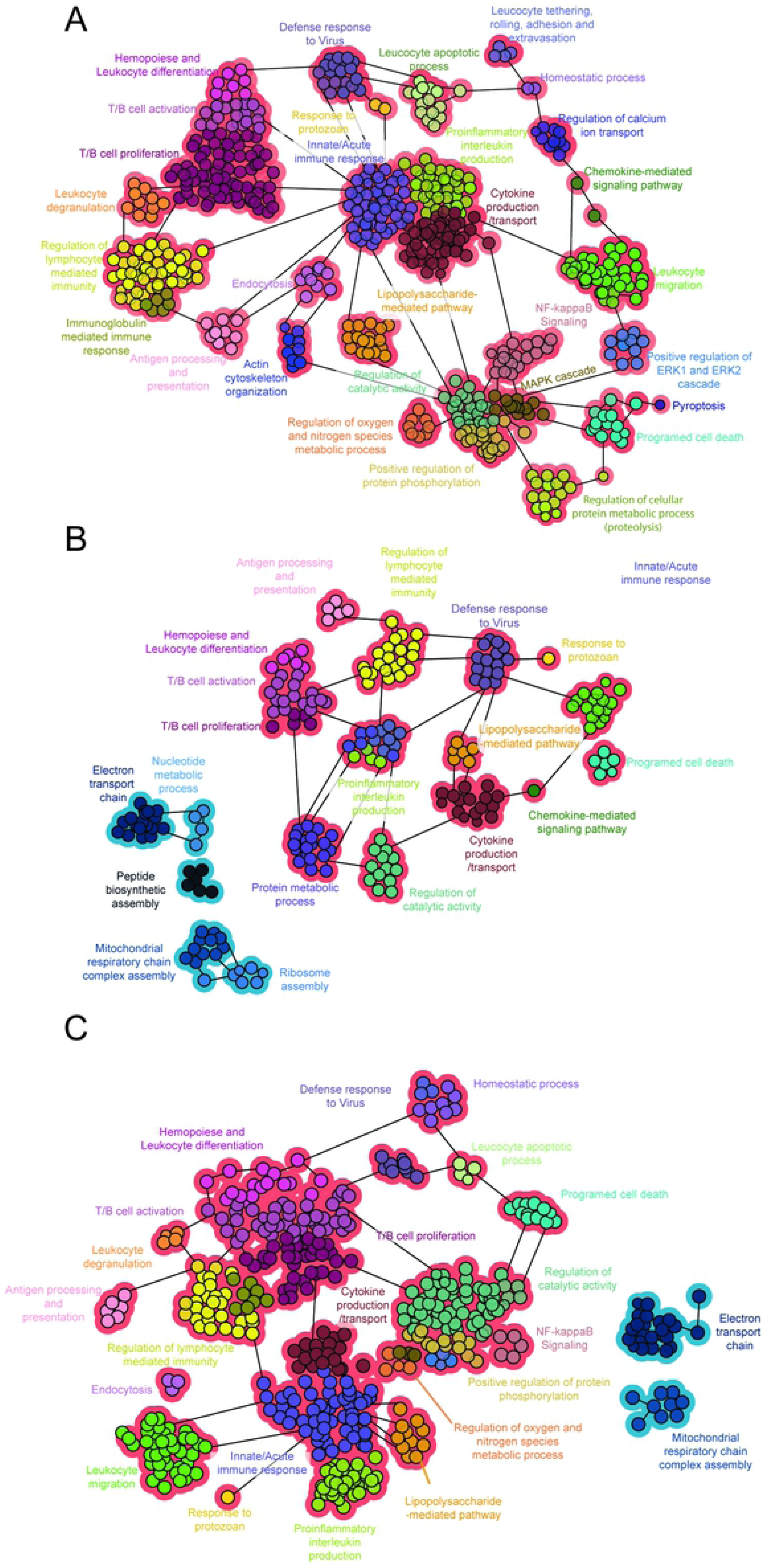
Functional networks of enriched biological processes. Network of biological processes of (A) Col1.7G2- (B) JG- and (C) Mixture-infected groups. Functional groups with more than 90% of upregulated or downregulated genes are shown in red or blue, respectively. Data represents only gene clusters with p-values less than 0.001 after Bonferroni step-down correction.

To visualize the effect of JG and Mixture-infected downmodulation of genes that play specific roles within the mitochondria, we generated a heatmap from the KEGG database. The complete expression data of all genes and experimental groups are presented in the supplementary material (S2 and S3 Table). Notably, from the 13 protein-coding genes of the mitochondrial genome, 7 genes (Nd1, Nd2, Nd4, Nd5, Nd6, Co1 and Cytb) were downregulated in all experimental groups. In the JG- and Mixture-infected animals, they were almost four-fold downregulated and were approximately two-fold downregulated in the Col1.7G2-infected group (S3 Table). In addition, large and small ribosome subunits were significantly downregulated in the JG-infected but not in the Col1.7G2-infected group (Fig 6A). Interestingly, metabolic pathways, such as Oxidative phosphorylation (Fig 6B) and Citric acid cycle were strongly downregulated in JG and Mixture-infected animals when compared to Col1.7G2. As previously noted, these findings suggests that JG promotes a drastic reduction in oxidative metabolism and protein synthesis in the infected cardiac cells. Previous microarray studies showed the same oxidative phosphorylation downregulation pattern in cardiomyocytes infected by other *T. cruzi* strains [40, 41]. The downregulation of the electron transport chain and oxidative phosphorylation gene expression could either increase or decrease reactive oxygen species (ROS) levels, depending on the proton motive force, NADH/NAD+ and CoQH_2_/CoQ ratios and O_2_ concentration [42]. However, previous studies have shown that ROS is a double-edged sword for *T. cruzi* parasites, as it acts as a signaling molecule for *T. cruzi* replication in macrophages at low concentrations [43, 44], but it can be harmful at higher concentrations [45]. Interestingly, treatment with catalase reduces the multiplication of JG in cardiomyocytes, but not Col1.7G2, suggesting that H_2_O_2_ acts as a signaling molecule for JG growth in these cells [36]. Notably, *T. cruzi* lacks the catalase gene, which may be important for the parasite to sense and overcome oxidative stress. Furthermore, *T. cruzi* parasites transfected with catalase showed increased resistance to H_2_O_2_ and impaired signals for cell differentiation compared with wild-type parasites [46]. Excessive ROS level causes oxidative DNA damage, such as 8-oxo-7,8-dihydroguanine (OG) accumulation, which slows pathogen growth. Remarkably, Campos et al. have shown that JG and Esmeraldo (both *T. cruzi II*) are more sensitive to oxidative DNA damage caused by H_2_O_2_ treatment when compared to Col1.7G2 and Silvio (both *T. cruzi I*) [47]. They also showed that even before treatment with 200 µM H_2_O_2_, JG displayed more oxidative lesions in DNA than Col1.7G2, as determined by the amount of OG present in the nucleus. The role of OG in *T. cruzi* replication signaling is still poorly understood, but recent studies have shown that parasites overexpressing MutT, which are responsible for removing OG from DNA, increased intracellular parasite replication (CL Brener strain) in fibroblasts compared to WT [45]. Interestingly, the 8-oxo-dGMP MutT product may act as a stress signal carrier, and OG DNA lesions might act as an epigenetic modification, which serves as a sensor for oxidative stress, increasing gene expression [48]. Thus, the available evidence suggests that the differential multiplication rate between different *T. cruzi* strains caused by sensing and responding distinctively to ROS, may be crucial to determine the strain colonization in the heart of mice, and possibly other mammals such as humans.

**Figure 6.**
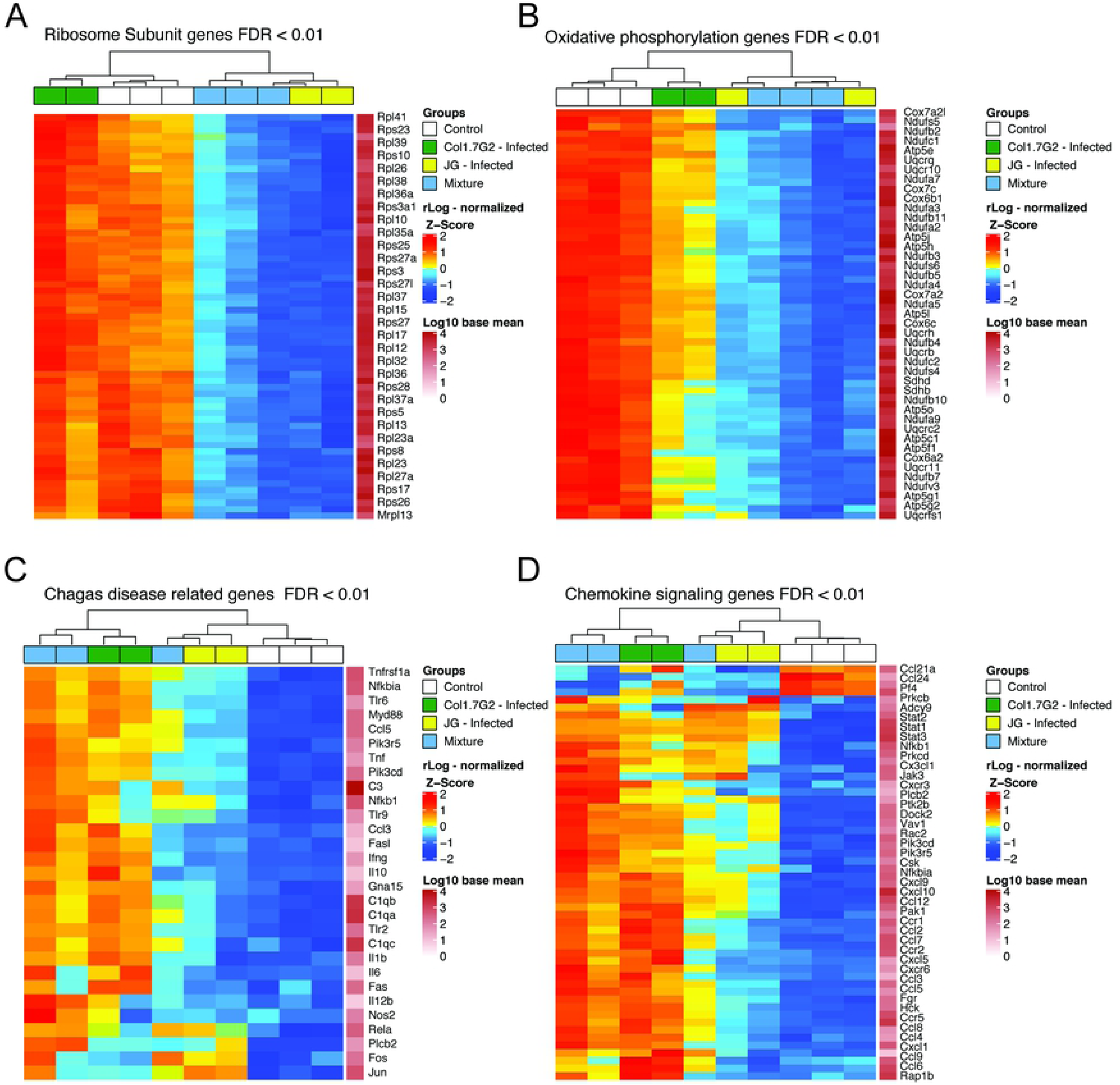
Heatmap of DEGs. included in (A) Ribosomes subunits, (B) Oxidative phosphorylation pathway, (C) Chagas disease related genes and (D) Chemokine signaling. Data represent z-score of normalized read counts (log2 scale) for each gene. Base mean track shows average gene expression between samples.

Next, we sought to visualize DEGs involved in the immune response detected as enriched biological processes to capture differences between *T. cruzi* strains during the acute stage of infection. To visualize DEGs involved with CD we accessed KEGG pathway 05142 which evidenced a clear distinction between the JG- and Col1.7G2-infected animals (Fig 6C). As noticed earlier, we observed that the JG-infected group exhibited weaker activation of several immune response genes compared to Col1.7G2. For instance, in Col1.7G2-infected animals, *Tlr9* and *Tlr2* had a positive fold-change of approximately 5, relative to control animals, while the JG-infected group showed a positive fold-change of 2.7 (Fig 6C, Table S3). The high expression profile of cytokines such as *Ifn-γ, Il-6, Tnf* and *IL-12* observed in Col1.7 G2-infected animals corroborates the previously described Th1 host response induced by *T. cruzi* [24]. However, JG-infected animals exhibit lower expression of these same genes. Similar to the CXC and CC chemokines involved in neutrophil and macrophage recruitment, transcriptional factors such as the Class II Major Histocompatibility Complex Transactivator (Ciita) and Tbx21/Tbet, a well-known Th1 cell-specific transcription factors, also displays lower expression levels in the JG group compared to the Col1.7G2 group (Fig 7C). Notably, animals inoculated with a mixture of both parasites generally exhibit the same pattern of immune response DEGs as the Col1.7G2-infected group. This expression pattern suggests that the Col1.7G2 strain is more likely to be detected early by the immune system and initiate a stronger immune response than the JG strain.

**Figure 7.**
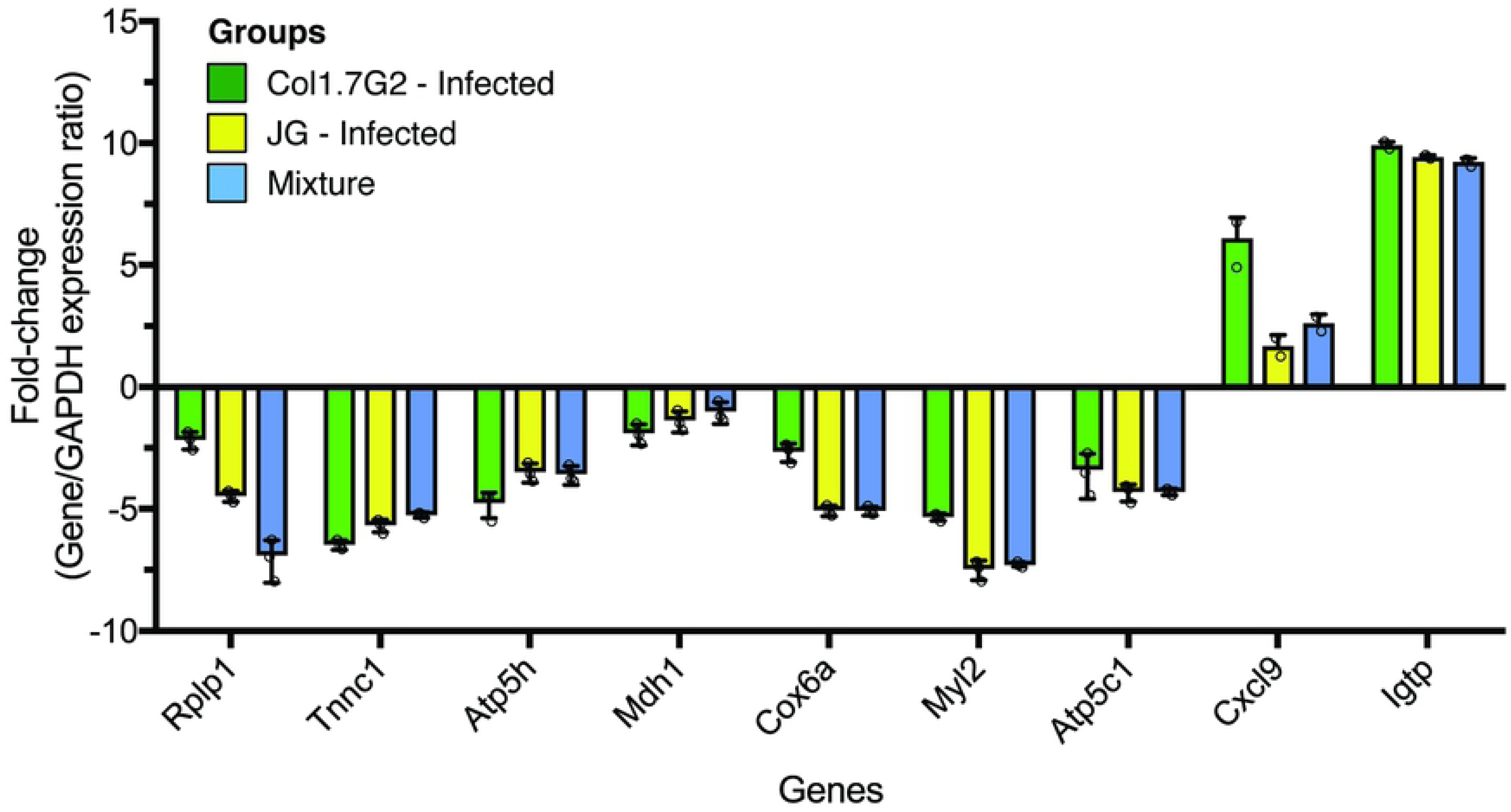
Quantitative real-time RT-PCR validation of differentially expressed genes. Expression levels of tested genes were normalized to Gapdh (housekeeping gene) and fold-change values were calculated with the 2^-ΔΔCT^ method. All PCR reactions were performed with three technical replicates.

The evasion capability of each strain can further explain why there are differences in the expression of genes involved in the immune response when animals are infected with different parasite strains. From earlier studies, parasite-specific CD8+ T cells were not seen in the bloodstream of BALB/c mice until day 9 of infection by the Y strain (a nonvirulent *T. cruzi II* strain similar to JG). To prove whether this effect was due to immune evasion or immunosuppression, the authors administered agonists of *Tlr-*2 and *Tlr-9* to emulate immune system recognition prior to infection. They showed that treated mice developed parasite-specific CD8+ T cells earlier than nontreated groups [49–51]. Therefore, the lower expression of immune response genes by JG-infected animals can be the result of an effective immune-system evasion of this strain, which could also be responsible for the persistence of this parasite in the heart of BALB/c mice in the chronic phase of the disease. One possible explanation for this evasion ability of *T. cruzi* is the transfer of sialic acid from host proteins to parasite surface proteins, such as mucins. Indeed, some of the most well-studied surface proteins of *T. cruzi*, namely, mucins, mucin-associated surface proteins (MASPs) and trans-sialidases (TSs) are strongly related to immune system evasion, cellular membrane adhesion and cellular invasion [52–56]. Nevertheless, *T. cruzi* I strains have fewer genes coding for the mucin, MASP and TS families compared to *T. cruzi II* [57].

To confirm the expression data obtained from the RNA-Seq, we performed qPCR for validation (Fig 7). Quantitative PCR data shows that the fold change of most tested genes corroborates the reported RNA-Seq expression tendency. Highly upregulated genes in RNA-Seq analysis, such as Cxcl9 and Igtp, are also seen to be upregulated by qPCR when compared to the Gapdh, and genes coding for proteins acting in the mitochondria are seen to be downregulated by both methods. Data represents one sample from each group with three technical replicates.

In the present work, we depicted the differential ability of two T. cruzi strains (JG and Col1.7G2) to modulate gene expression in hearts of BALB/c mice at the acute phase of infection. We also showed how the mixture of these *T. cruzi* strains affects host gene expression. We described two major distinct behaviors: hearts of mice inoculated with JG or with the mixture of both strains exhibit many genes from oxidative metabolism and translation downregulated compared to the uninfected controls. However, hearts of mice inoculated with Col1.7G2 and with the mixture of both strains showed a strong activation of genes from the innate and adaptive immune response compared to noninfected hearts. Corroborating previous findings, we propose here that the remarkable differences between the two *T. cruzi* strains in their ability to persist in BALB/c hearts in the chronic stage of CD could be explained by differences in the higher intracellular proliferation rate of JG and its ability to slowly activate the immune response of the host at the acute phase of infection, in contrast to Col1.7G2, which strongly activates the host immune response and has a slower proliferation rate in cardiomyocytes. JG parasites would also benefit from the robust reduction of the overall energetic status of the cell and augmentation of ROS production to better establish in the tissue (Fig. 8). We suggest here that the different features of each *T. cruzi* strain, such as ROS signaling, proliferation and immune system evasion, could determine the survival of one strain over others and prevail in host tissues. Altogether, our research highlights the need for a better understanding of the effect of *T. cruzi* polyparasitism, as well as the uniqueness of each *T. cruzi* strain and its interaction with the host.

**Figure 8.**
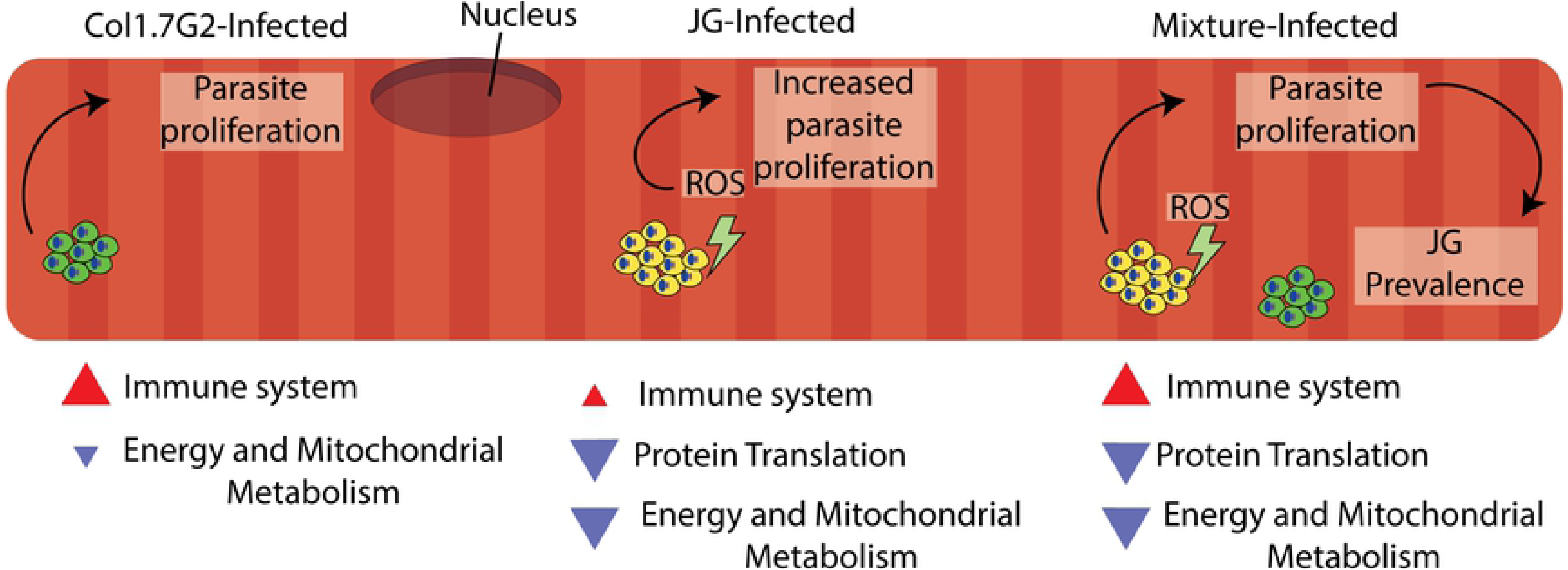
Proposed model for preferential prevalence of JG over Col1.7G2 in BALB/c mouse hearts. Throughout the acute stage of *T. cruzi* infection, Col1.7G2 parasites induce a strong Th1-polarized immune response in mouse hearts. Although a potent Th1 response helps developing against parasites, amastigote nests are capable to survive later at chronic stages. Mice infected with the JG strain exhibit lower Th1 response and higher survival rates compared to Col1.7G2. In parallel, JG parasites, stimulated by higher sensitivity to ROS generated by mitochondrial metabolism down-modulation, display rapid increase in clonal burden. During a mixed infection with both strains, JG parasites can overcome Col1.7G2 population due to an enhanced proliferation.

## Materials and Methods

### Ethics Statement

All procedures complied with the standards stated in the Guide for the Care and Use of Laboratory Animals and were conducted under conditions approved by the local animal ethics committee [58]. The Institutional Committee for Animal Ethics of UFMG (CEUA-UFMG, license 64/12) approved all experimental procedures used in this study. Both noninfected and infected animals were kept in plastic boxes with food and water *ad libitum* with the same appropriate conditions of technical management in cages that were properly identified and sealed in a 12-h light dark cycle environment. All animal procedures were performed under anesthesia using a mixture of ketamine (50 mg/kg) and xylazine (10 mg/kg).

T. cruzi populations used in this study, Col1.7G2 and JG, belong to lineages TCI and TCII, respectively. JG strain was originally isolated in 1995 by Professor Eliane Lages-Silva (Universiade Federal do Triangulo Mineiro, Brazil) from a chronic patient with megaesophagus. Col.1.7G2 strain is a clone from Colombian strain, which was originally isolated by Federici in 1964 from a chronic patient with cardiac disorder [59]. Both JG and Col1.7G2 were formerly characterized as monoclonal population, through the analysis of the eight microsatellite loci according to previously described methodology [60]. Both strains belong to the *T. cruzi* collection from Laboratory of Chagas Disease (UFMG), coordinated by Professor Egler Chiari. Infective trypomastigote forms were thawed from liquid nitrogen preservation and parasites were injected into SWISS mice for population expansion and diluted to 50 parasites/100 µl of sterile PBS for further use in BALB/c infections.

### Experimental Infection

Inbred 6-8-week-old male BALB/c mice were obtained from the Centro de Bioterismo (CEBIO/ICB, Belo Horizonte, Brazil) and housed in our local animal facility under the same conditions. Mice were randomly divided into four groups of 5 individuals each. The noninfected controls were injected intraperitoneally (i.p.) with phosphate-buffered saline (PBS-vehicle), while the three experimental groups were injected (i.p.) with 50 trypomastigotes of the JG (*T. cruzi* II), 50 of Col1.7G2 (*T. cruzi* I) and an artificial mixture of both strains (50 + 50 parasites). After infection, mice were caged according to group. Blood parasitemia was verified by counting the parasites present in 5 µL of blood using an optical microscope at the 14^th^ day post infection. On the 15^th^ day, mice were anesthetized, and their hearts were collected, rapidly washed in sterile PBS and immediately frozen in liquid nitrogen.

### RNA extraction, quality assessment, and sequencing

Frozen hearts were kept in liquid nitrogen and thoroughly pulverized into a powder using a sterile porcelain crucible. The total RNA for sequencing was extracted using the Trizol reagent (Life Technologies) following the manufacturer’s protocol and precipitated with isopropanol. The quality and integrity of the samples were verified by electrophoresis on a Bioanalyser 2100 (Agilent Technologies), and only samples with an RNA integrity number (RIN) greater than seven were selected for library construction (S9 Table).

The cDNA libraries were prepared and sequenced at the Beijing Genomics Institute (Shenzhen, China). In short, polyadenylated RNA was purified from total RNA, converted to cDNA using random hexamer primers, sheared, and size-selected for fragments ∼200 bp in length using the Illumina TruSeq RNA Sample Preparation Kit v2. Sequencing was performed on the Illumina Hiseq 2000 (Illumina, CA) platform and generated approximately 12 million paired-end reads, which were 90 nucleotides in length for each sample. Each group was sequenced in triplicate.

### Processing of raw sequencing reads

Raw reads were first checked for quality using FastQC (Babraham Bioinformatics, Cambridge, UK) [61]. Since all samples displayed high-quality scores, no sequence trimming was performed. STAR aligner (v2.5) was used with default parameters to map read locations in the mm10 (GRCM38) mouse reference genome [62]. Only uniquely mapping reads were retained for use in the downstream analysis (S1 Fig).

### Differential Expression, and graphic visualization

Information on transcript abundances was obtained by raw read counts using HTseq-count [63], and differential expression analysis was performed using DESeq2 [64] in R version 3.2.4. For this study, we considered as differentially expressed genes presenting a 2-fold change in relation to the control group, with a false discovery rate (FDR) of 1%.

Principal component analysis (PCA) was performed using the regularized-logarithm transformation (rlog) offered by the DESeq2, and the heatmaps were constructed using the heatmap.2 function of gplots. A Venn diagram was created with an area-proportional web application [65].

All figures were mounted and treated using vector graphics quality within Adobe Illustrator 2017.

### Enrichment analysis

We conducted the enrichment analysis using an R package with mouse genome-wide annotations [66]. Functional enrichment for biological processes (based on implemented Fisher’s exact test) was calculated with the Bioconductor package topGO [67]. Of note, topGO makes a nonbiased background adjustment of data for enrichment analysis, since it considers as background only non-DE genes with similar expression to DE genes as controls for comparison. We model the background by considering only genes expressed in the samples analyzed. In addition, we analyzed the upregulated and downregulated genes separately, as described previously [68].

### Functional network analysis

The networks of enriched functional terms were constructed with the ClueGO plug-in using Cytoscape software [69, 70]. Our lists of DEGs was analyzed using the biological process EBI-QuickGO mouse annotation. Pathways with p-values less than 0.001 were considered statistically significant after Bonferroni step-down correction.

### Quantitative Real-Time PCR (qPCR)

Total RNA of mouse hearts was extracted using TRIzol (Life Technologies, CA, US) and treated with TURBO^TM^ DNase (Thermo Fisher Scientific, MA, US). cDNA was then produced with Superscript II kit first strand synthesis (Invitrogen, CA, US). PCR experiments were performed in a 7900HT Fast Real-Time PCR System (Applied Biosystems, CA, US) using the PowerUp SYBR Green Master Mix (Thermo Fisher Scientific, MA, US). Tested genes and primer sequences are provided in the Supplementary Table 10.

### Data availability

All raw and processed sequencing data are available in the GEO (Gene Expression Omnibus) under the accession code: GSEXXXXXX.

## Acknowledgments

The authors would like to thank the valuable contribution of Neuza Antunes Rodrigues and Afonso da Costa Viana who assisted us with mice procedures and parasite handling and Fabricio Rodrigues dos Santos for making the use of the 7900HT Fast Real-Time PCR System available for qPCR analyses.

## Author Contributions

**Conceptualization:** Castro TBR, Franco GR, Macedo AM, Canesso MCC, Pena SD

**Data Curation:** Castro TBR, Chame DF, de Laet Souza D

**Formal Analysis:** Castro TBR

**Funding Acquisition:** Macedo AM, Franco GR, Chiari E, Machado CR, Pena SD, Castro TBR

**Investigation:** Castro TBR, Chame DF, Laet Souza D

**Methodology:** Castro TBR, Franco GR, Macedo AM, Toledo NE, Boroni M

**Project Administration:** Franco GR, Macedo AM

**Resources:** Chiari E, Macedo AM, Franco GR, Pena SD, Machado CR, Tahara EB

**Supervision:** Franco GR, Macedo AM

**Validation:** Castro TBR, Chame DF, de Laet Souza D

**Visualization:** Castro TBR

**Writing – Original Draft Preparation:** Castro TBR, Franco GR, Macedo AM, Canesso MCC, Boroni M, Toledo NE, Chame DF, Laet Souza D, Tahara EB, Pena SD

## Supporting information

**S1 Table. Total number of reads and mapping statistics.** Number of sequences from each sample and proportion of reads mapping at single positions using GRCm38 mouse genome.

**S2 Table. Dataset of all detected genes.** Full dataset of gene expression analysis after comparing experimental groups with controls, including Ensembl gene ID, external gene name, gene biotype, chromosome coordinates, expression mean, fold change statistics and adjusted p-values (FDR).

**S3 Table. Filtered dataset from genes considered differentially expressed.** Complete list of differentially expressed genes (DEG) from all experimental groups after filtering the full dataset of detected genes following a 2-fold change and 1% FDR.

**S4 Table. Gene biotype statistics.** Statistics of all differentially expressed genes with their biotype classification.

**S5 Table. Venn diagram dataset.** List of all shared genes in each experimental group and their expression levels which were used to generate the Venn diagram.

**S6 Table. Complete Gene Ontology analysis dataset.** Table with a complete list of terms containing up- or downregulated genes. Each functional category is shown with the number of detected (annotated) and differentially expressed (significant) genes.

**S7 Table. Differentially expressed genes per GO category.** Table with a complete list of all DEGs encountered in each enriched GO category.

**S8 Table. Genes per GO category.** Table with a complete list of all expressed gene encountered in dataset per enriched GO category.

**S9 Table. Sample description and RNA quality statistics.** RNA quality attributes of samples used in library construction.

**S1 Fig. Overall workflow.** All major bioinformatics steps performed in this work.

